# Microscale nonrandomness revisited: An overlooked predictor of ecological change

**DOI:** 10.64898/2026.11.05.764087

**Authors:** Takeshi Miki, Kinuyo Yoneya, Ayumi Katayama, Si Ho Han, Housei Nakamura, Shungo Takeda, Hiroto Miyawaki, Tomonori Kume, Kanako Shirotsuka, Shuji Kaneko, Kensuke Seto, Silke Van den Wyngaert, Maiko Kagami

## Abstract

Abrupt ecological transitions have driven the development of early warning signal metrics that track how ecological systems respond as external conditions gradually change and systems approach transitions between states. However, their generality and applicability remain uncertain, particularly for short-term ecological dynamics that can occur without directional external change. To address this gap, we propose a general framework based on the hypothesis that microscale nonrandomness, quantified from frequency distributions or spatial patterns, predicts subsequent short-term rates of change in populations and interactions, as well as current macroscale ecosystem states. We applied the framework to four datasets: time series from an incubation experiment of phytoplankton–parasitic chytrid interactions; spatiotemporal field observations of leaf beetles on willow trees and aphids in a crop system; and spatial snapshot data linking soil surface characteristics with aboveground and soil ecosystem properties in a forest vegetations–soil system. We quantified microscale nonrandomness using Morisita I_δ_ from frequency distributions and Moran’s I and SADIE from spatial patterns. Across all systems, models incorporating microscale nonrandomness metrics consistently showed greater cross-validated predictive performance than those based on macroscale variables or spatial averages alone. Although their relative performance varied among systems, frequency-based metrics performed as well as or better than spatially explicit metrics in some cases. The consistent improvement across contrasting systems supports the broad applicability of this framework, including under limited data conditions. This framework broadens ecological forecasting to short-term, continuous dynamics beyond those associated with directional environmental change.

**Significance statement:** Ecological forecasting has largely focused on major shifts in ecosystem states, often associated with gradual changes in environmental conditions. However, ecosystems also undergo short-term, continuous changes that are difficult to predict using existing approaches. Here, we demonstrate that microscale nonrandomness in the distribution of organisms and environmental properties can improve predictions of ecological rates in the near future, such as population growth and changes in prevalence, as well as current ecosystem states. Consistent improvements across four contrasting ecological systems highlight the broad applicability of this approach, even with limited data. Our findings expand the scope of ecological forecasting by revealing the predictive value of fine-scale ecological patterns.

## Introduction

Abrupt transitions between alternative stable states (ASS) have been an important focus in ecological forecasting, particularly because they represent critical changes with substantial ecological and societal consequences. Consistent with this focus, well-documented examples of ASS occur in lakes (Carpenter, Ludwig & Brock, 1999; Su *et al*., 2021), terrestrial vegetations (Rietkerk *et al*., 2004), and coral reefs (Mumby, Hastings & Edwards, 2007). These observations have motivated the development of early warning signal (EWS) metrics for detecting or anticipating rapid regime shifts (Scheffer *et al*., 2009; Dakos *et al*., 2015; Grziwotz *et al*., 2023). Many of these metrics are based on bifurcation theory, including approaches to detect critical slowing down (CSD: Kuehn, 2011) and methods for dynamically estimating dominant eigenvalue (Grziwotz *et al*., 2023). Their performance has been demonstrated across multiple disciplines, including ecological, biomedical, and climate sciences (Dakos *et al*., 2024).

However, several considerations suggest that focusing ecological forecasting primarily on such abrupt transitions, and on the assumptions underlying EWS-based approaches, may be more context-dependent than often assumed.

First, key assumptions underlying the EWS framework are not always satisfied across ecological systems. Although ASS have been documented in a range of ecosystems, their generality remains debated. For example, evidence for ASS is limited or inconclusive in some coral reef systems (Żychaluk *et al*., 2012), and meta-analyses have questioned their ubiquity across ecosystems (Hillebrand *et al*., 2020). In addition, the statistical power to detect ASS is not always high (Hillebrand *et al*., 2020). Theoretical studies further suggest that when additional ecological complexity is incorporated into models already characterized by strong positive feedback loops, e.g., plant–soil feedback mediated by microbial diversity, coral–algal systems with physical legacy effects, and canonical models with stochasticity and spatial heterogeneity, these feedbacks can be weakened or offset by additional processes, thereby reducing the conditions under which ASS arise (Miki *et al*., 2010; Cheung, Nozawa & Miki, 2021; Das, Shnerb & DeMalach, 2026). Empirical analyses of lake ecosystems indicate that transitions driven by stability shifts between ASS represent only a subset of observed ecological changes (O’Brien *et al*., 2023). Moreover, some population models incorporating nonlinear dynamics exhibit abrupt transitions without ASS (Hastings & Wysham, 2010). Together, these findings suggest that both the prevalence of ASS and the role of bifurcation-driven stability loss in ecological transitions may be more limited or context-dependent than often assumed.

Second, even when these assumptions are satisfied, the performance of EWS-based approaches can be variable. For example, even in transitions governed by bifurcation-driven stability loss, some EWS metrics do not consistently detect such transitions (O’Brien *et al*., 2023). While certain indicators, particularly those based on CSD (e.g., variance and autocorrelation), may have broader applicability (Dakos *et al*., 2015), their performance can still vary depending on system characteristics and data structure (Weinans *et al*., 2021).

Beyond these considerations, there are also practical and conceptual limitations to current EWS approaches. Many EWS approaches require long-term, high-frequency time series (Weinans *et al*., 2021) or large-scale spatial monitoring (Dakos *et al*., 2010; Kéfi *et al*., 2014), which can limit their applicability in systems where only short time series or snapshot observations are available, such as in small-scale agriculture, forestry, and fisheries (Pauly & Zeller, 2016; Jiménez *et al*., 2019; de Groot *et al*., 2023). In addition, EWS approaches have primarily been developed to detect gradual or abrupt shifts in system states over relatively long timescales, often at the inter-annual scale. However, ecological change also includes shorter-term, continuous variations in system variables, such as fluctuations in population growth rates or interaction strengths, which can occur even in the absence of directional changes in external drivers (Hastings, 2004; Kidd & Amarasekare, 2012). In applied contexts such as integrated pest management, degree-day models have successfully predicted the timing of key events, including pest emergence or peak abundance (Herms, 2004; Kaneko, Shirotsuka & Shibao, 2017; Day *et al*., 2024), but such approaches are typically system-specific and do not aim to forecast the whole trajectory of population dynamics. As a result, these short-term dynamics remain less well captured by more general approaches and may require alternative types of metrics that do not rely on extensive temporal or spatial data. Such variability may also be associated with shifts in microscale aggregation or heterogeneity that are not detectable from macroscale state variables alone (Favis-Mortlock, 1998; Holfeld, 2000; Lloyd-Smith *et al*., 2005; Stein, 2011; Yoneya *et al*., 2021).

To address these limitations, we approach ecological prediction from a different perspective, focusing on microscale nonrandomness as a potential predictor of short-term ecological dynamics and current ecosystem states, without relying on assumptions about alternative stable states. We hypothesize that microscale nonrandomness, whether spatially explicit or frequency-based, reflects shifts in aggregation or heterogeneity that precede or accompany changes in macroscale variables. This hypothesis is motivated in part by studies of spatial EWS metrics, in which self-organized patterns have been interpreted as footprints of local feedback processes (Rietkerk *et al*., 2004). Accordingly, shifts in microscale patterns may serve as predictors of (i) subsequent rate changes when temporal data are available, or (ii) macroscale state variables when only snapshot data are available.

We translate this idea into a predictive framework using established metrics of nonrandomness, including frequency-based (spatially implicit) indices such as Morisita I_δ_ (Morisita, 1962; Hurlbert, 1990; Yoneya *et al*., 2021; De Freitas Alves & De Santana, 2022) and spatially explicit indices such as Moran’s I (Moran, 1950; Legendre & Fortin, 1989) and SADIE (Perry, 1995, 1999; Li, Madden & Xu, 2012; Winder *et al*., 2019). Although Moran’s I has been applied in spatial EWS studies to quantify spatial autocorrelation (Dakos *et al*., 2010), and SADIE has long been used to describe aggregation patterns (Turechek & Madden, 1999; Winder *et al*., 2001; Maestre & Cortina, 2002), their use as predictors of macroscale ecological dynamics or states has remained limited. Unlike standard EWS approaches, this framework does not require long-term time series but instead relies on high-resolution observations at finer scales, enabling application across systems with differing data availability.

We evaluate the generality of this approach across ecological contexts that differ in system type and data structure. For rate-based analyses, we consider resource–consumer systems in both aquatic and terrestrial settings, including host–parasite interactions in a freshwater lake and plant–insect interactions (tree–herbivore and crop–pest systems), using temporal data with frequency-based (spatially implicit) metrics and, where available, spatially explicit metrics. For state-based analysis, we examine vegetation–soil interactions in a forest using snapshot data with spatially explicit metrics. These systems have ecological and practical relevance, as forecasting their population dynamics or assessing their current states can contribute to understanding ecological change and, in some cases, ecosystem or resource management (see **SI Materials and Methods** for the ecological context of each system). Together, these case studies differ in distinct ecosystem types, interaction structures, and data availability, providing a basis to assess the generality of the proposed framework.

## Results

For rate-based analyses, we developed generalized least-squares models with an autoregressive correlation structure to forecast the specific growth rate of focal variables from time step *k* to *k* + 1 using macro- and microscale state variables at time step *k*. First, in the diatom host–chytrid parasite system, we inoculated the growth medium with low initial abundances of the diatom and chytrid on day 0 and monitored their dynamics until day 10. The four replicate units exhibited substantial temporal variation in host abundance, parasite abundance, and parasite prevalence **(Fig. 1A**). We then focused on forecasting the specific growth rates of parasite abundance and prevalence from these time-series data. The distribution of infection intensity, i.e., the number of sporangia per host cell (e.g. *SI Appendix* **Fig. S1**), often deviated from a Poisson distribution (**Fig. 2A**; Yoneya et al. 2021). The Morisita I_δ_ index, a microscale frequency-based (spatially implicit) metric of nonrandomness, was retained in the best performing models for both parasite abundance and prevalence in cross-validation forecasting (*SI Appendix* **Table S1**). Its contribution to forecasting performance varied among replicate units but was substantial, particularly for prevalence, compared with models using macroscale variables alone (**Fig. 3AB**). One-step-ahead abundance forecasts, reconstructed from observed abundance and forecasted growth rates for a representative replicate unit, further indicated improved forecasting of abundance changes when the Morisita index was included (**Fig. 4AB**).

**Figure 1.**
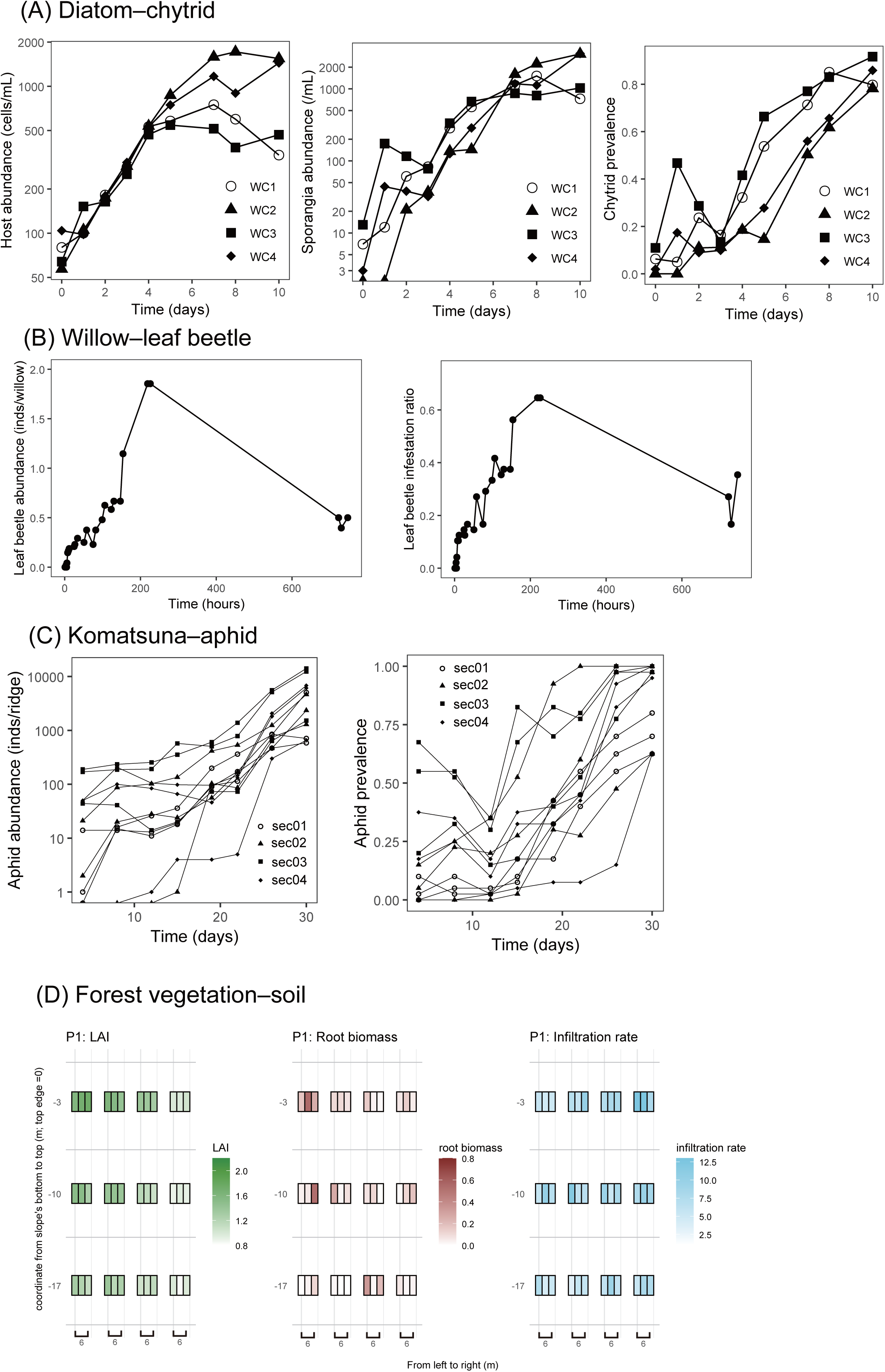
Target variables for forecasting and prediction across four ecological systems. (A) Diatom– chytrid system: time series of host abundance, chytrid sporangia abundance, and chytrid prevalence. (B) Willow–leaf beetle system: time series of leaf beetle abundance and leaf infestation rate. (C) Komatsuna–aphid system: time series of aphid abundance and aphid prevalence. (D) Forest vegetation–soil system: spatial distributions of leaf area index (LAI), root biomass, and soil infiltration rate.

**Figure 2.**
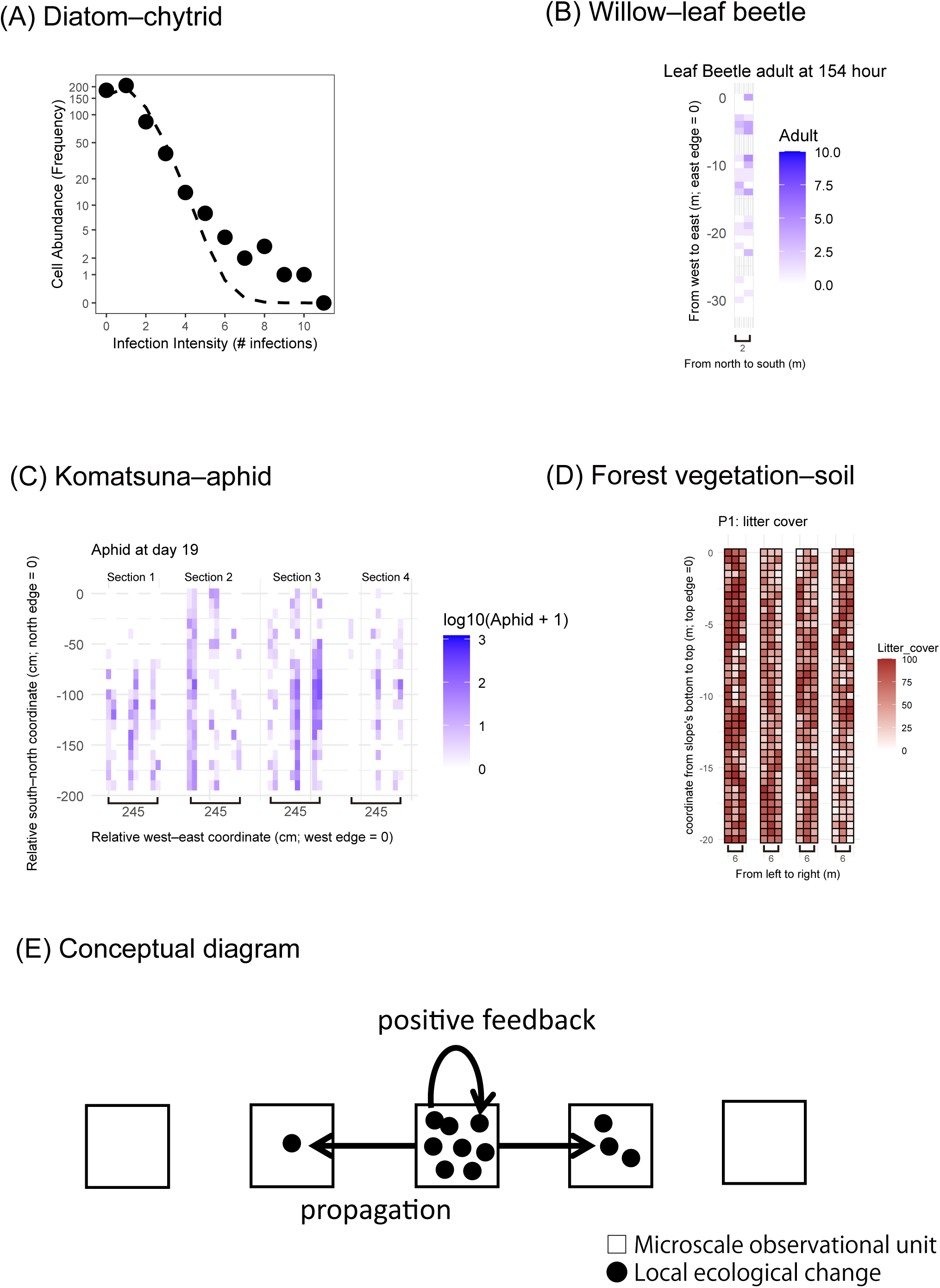
Microscale patterns quantified across four ecological systems. (A) Diatom–chytrid system: relationship between infection intensity and host cell abundance. The dashed line indicates the expected relationship under a random distribution (i.e., Poisson distribution) of infections among host cells. (B) Willow– leaf beetle system: representative spatial distribution of adult leaf beetles at 154 h. (C) Komatsuna–aphid system: representative spatial distribution of aphid abundance at day 19. (D) Forest vegetation–soil system: representative spatial distribution of litter cover, one of eight microscale soil-surface variables measured at each census point. Spatial distributions of all eight variables are shown in *SI Appendix* **Fig. S4**. (E) Conceptual diagram illustrating how initially small local changes may be amplified by positive feedback and subsequently propagate among biological units or through physical space, generating microscale nonrandomness. Observational units can represent discrete biological units, such as individual host cells, or units defined in physical space.

**Figure 3.**
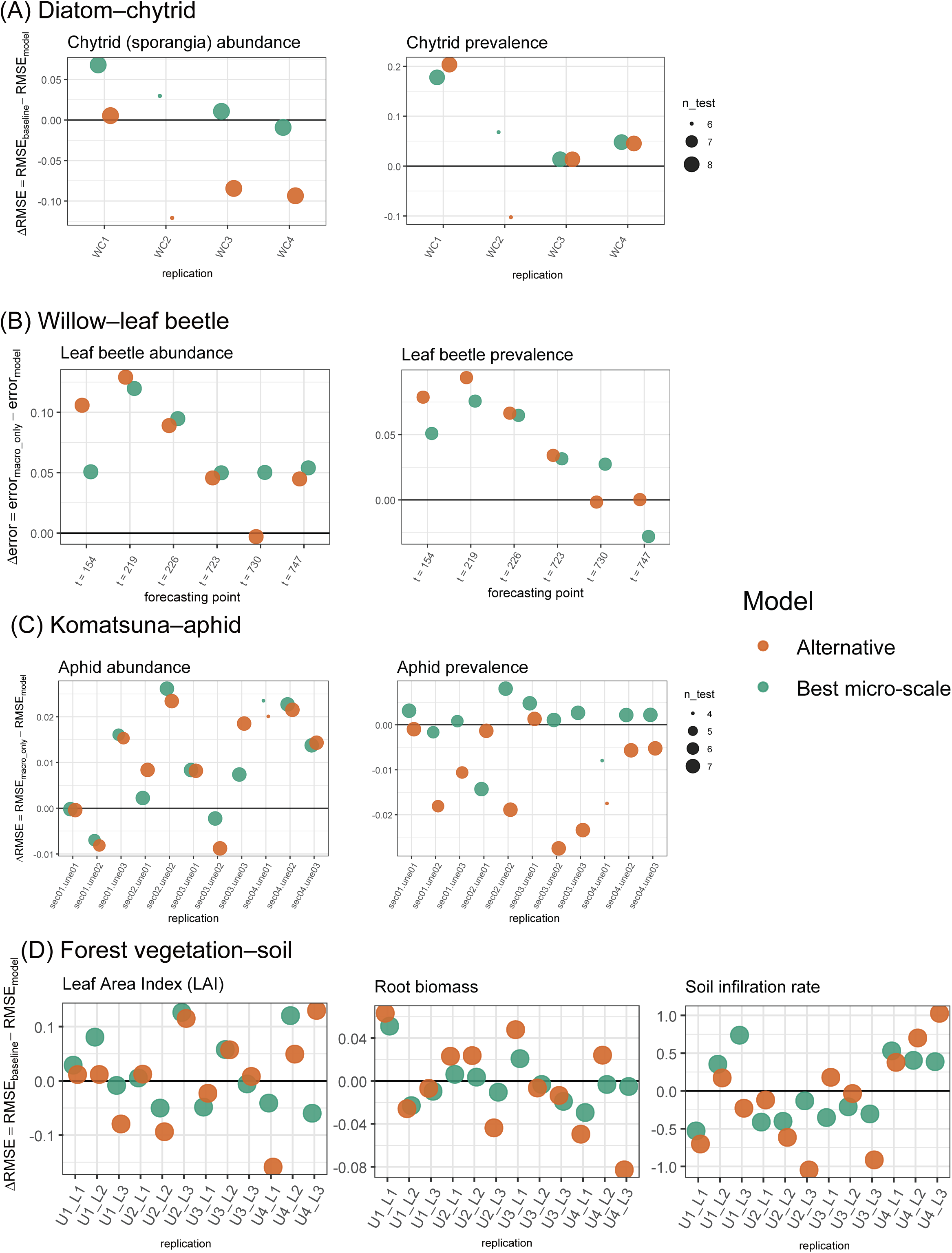
Forecasting and prediction performance of models incorporating microscale metrics relative to baseline models. Performance differences between models incorporating microscale metrics and the corresponding baseline models are shown for individual evaluation units in the (A) diatom–chytrid, (B) willow– leaf beetle, (C) Komatsuna–aphid, and (D) forest vegetation–soil systems. For each target variable, the baseline was the best-performing model among those without microscale metrics. For the diatom–chytrid, Komatsuna– aphid, and forest vegetation–soil systems, ΔRMSE was calculated as RMSE_baseline_ – RMSE_model_ for each held-out replicate or unit. For the willow–leaf beetle system, in which each forecasting point provided a single test observation, ΔAE was calculated as AE _baseline_ − AE_model_, where AE is absolute prediction error. Thus, positive values indicate improved forecasting or prediction relative to the baseline model. Green points represent the best-performing model among those incorporating microscale metrics; these were also the overall best-performing models for all target variables except root biomass, for which the baseline model performed best overall. Orange points represent alternative models incorporating microscale metrics. Point size indicates the number of observations used to evaluate prediction error. Detailed model structures are provided in *SI Appendix* **Tables S1-S4**.

**Figure 4.**
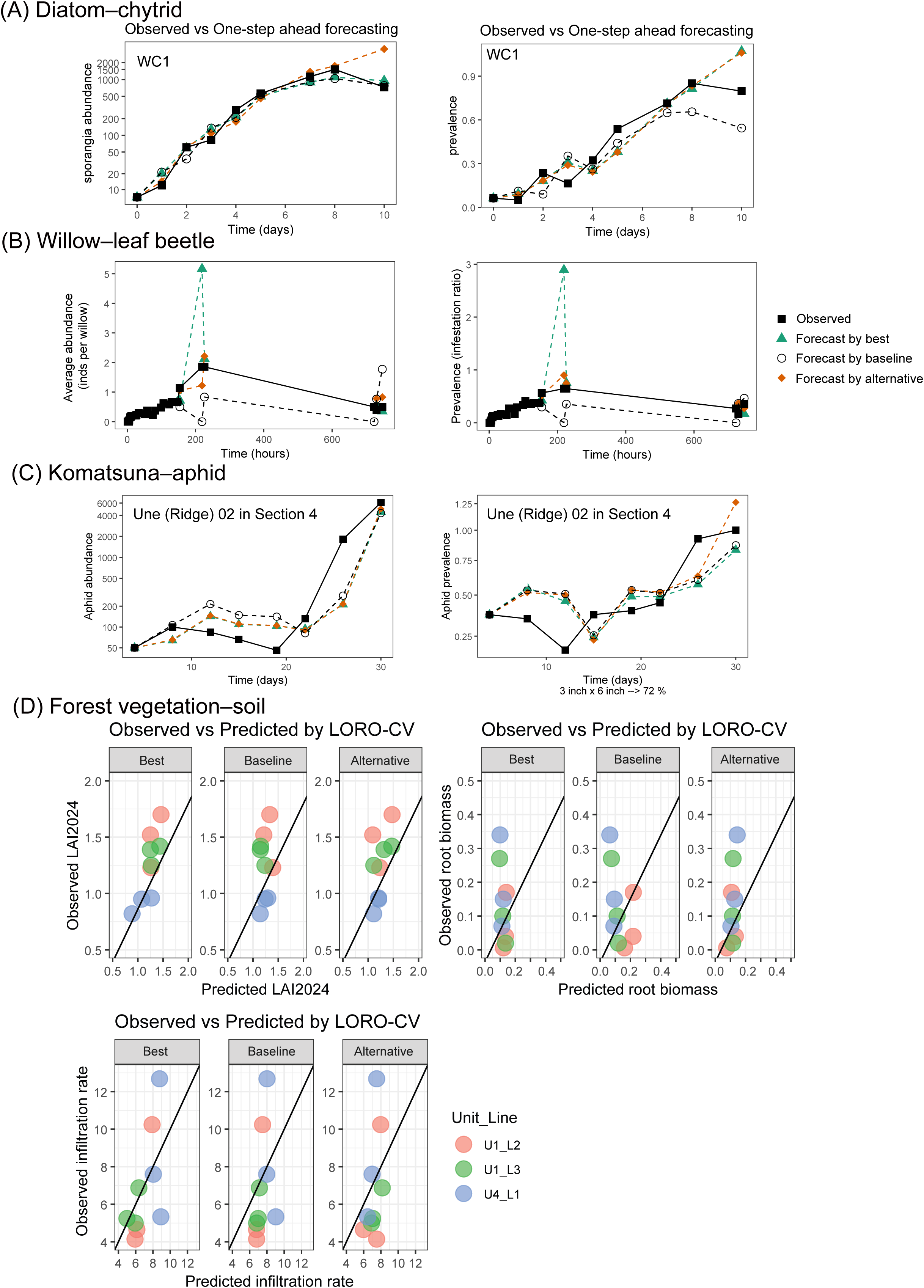
Illustration of forecasts and predictions from models incorporating microscale metrics. (A–C) Observed state variables and one-step-ahead forecasts for the (A) diatom–chytrid, (B) willow–leaf beetle, and (C) Komatsuna–aphid systems. The models forecasted growth rates, and their performance was evaluated from errors between observed and forecasted growth rates (Fig. 3), rather than from the state-variable trajectories shown here. For visualization, each one-step-ahead forecast shown here was calculated from the observed state variable at the preceding time point and the growth rate forecasted by the corresponding model. Green and orange lines indicate forecasts from the best-performing and alternative models incorporating microscale metrics, respectively, and black dashed lines indicate forecasts from the baseline models. At 723 h in the willow– leaf beetle system, forecasts from the best-performing model and those from the alternative models are omitted because they greatly exceed the plotted ranges (leaf beetle abundance, 1.60×10^4^ & 1.30×10^5^; prevalence, 1.97×10^3^ & 5.41×10^3^). (D) Forest vegetation–soil system: observed versus predicted leaf area index (LAI), root biomass, and soil infiltration rate under leave-one-replicate-out cross-validation (LORO-CV). Unlike A–C, these state variables were themselves the prediction targets, and their prediction errors were used directly to evaluate model performance in Fig. 3D. Diagonal lines indicate 1:1 relationships between observed and predicted values. For each target variable, the baseline was the best-performing model among those without microscale metrics. For root biomass, the baseline was the overall best-performing model, whereas the model labeled “Best” was the best-performing model among those incorporating microscale metrics.

Second, in the willow–leaf beetle system, we tracked the early phase of population dynamics following free immigration of leaf beetles onto intact willow individuals, with frequent observations during the first 1–12 hours and over subsequent days up to day 32 (**Fig. 1B)**(Yoneya, Miki & Takabayashi, 2023). We then focused on forecasting the specific growth rates of leaf beetle abundance and prevalence from these time-series data. From leaf beetle abundance on each willow individual at each time point (48 willow individuals observed across 27 time points; **Fig. 2B**; *SI Appendix*. **Fig. S2**), we quantified the standardized Morisita index as a frequency-based (spatially implicit) metric, and Moran’s I and the LMX index as spatially explicit metrics (**Figs. S2, S3**). Because the dataset consisted of a single time series, forecasting performance was evaluated using one-step-ahead forecasting rather than cross-validation. Models including these microscale metrics outperformed those using the macroscale variable alone (prevalence) for forecasting the growth rates of both abundance and prevalence (**Figs. 3CD, 4CD**, and *SI Appendix* **Fig. S4**), and the best forecasting models for both response variables retained only microscale metrics (Morisita index), without any macroscale variables (*SI Appendix* **Table S2**).

Third, in the Komatsuna–aphid system, we similarly focused on the early phase of aphid population dynamics following artificial colonization of aphids onto intact crop individuals under replicated experimental units (**Fig. 1C**). We then focused on forecasting the specific growth rates of aphid abundance and prevalence from these time-series data. From aphid abundance on each plant individual (480 Komatsuna individuals in total: 40 individuals per replication unit (ridge) × 3 units per section × 4 sections, observed across 8 time points over 30 days; **Fig. 2C**; *SI Appendix* **Fig. S2**), we quantified the standardized Morisita index as a frequency-based (spatially implicit) metric, and Moran’s I and the LMX index as spatially explicit metrics (*SI Appendix* **Fig. S3**). Models including these microscale nonrandomness metrics outperformed those using macroscale variables alone in cross-validation forecasting (**Figs. 3, 4**, and *SI Appendix* **Fig**. **S4**). The best forecasting model for aphid abundance growth retained population size together with the standardized Morisita index and the LMX index, whereas that for prevalence growth retained prevalence together with the LMX index (*SI Appendix* **Table S3)**. Forecasting performance remained partly robust even when both macroscale variables (population size and prevalence) and microscale metrics (Morisita and LMX) were calculated from only half of the original dataset, while the growth rates to be forecasted were calculated from the full dataset (*SI Appendix* **Fig. S2D**). Under this reduced-data condition, the best forecasting model for aphid abundance growth retained only the Morisita index, whereas that for prevalence growth retained only macroscale variable (i.e. prevalence) calculated from the halved dataset (*SI Appendix* **Table S3**).

Finally, for the state-based analysis of vegetation–soil interactions, we aimed to predict three macro-ecological variables: LAI, representing above-ground biological function; root biomass, representing a below-ground biological functional proxy; and soil infiltration rate, representing physical soil function. These variables were measured at three points (3, 10, and 17 m) along each 20 m line transect (36 sampling points in total: 3 points per transect × 3 transects per unit × 4 units; **Fig. 1D**). As predictors, we quantified eight microscale soil surface characteristics, including the presence or absence of pillars, cover by litter, vegetation, and stones, and litter layer thickness, within 0.2 × 0.2 m observation windows placed every 0.5 m along the same transects (41 points per transect × 3 transects per unit × 4 units; **Fig. 2D**, *SI Appendix* **Fig.S5**). For each macroscale sampling point, we calculated spatial averages of the eight microscale variables and their corresponding LMX indices and Moran’s I values. Each set of eight metrics was then represented by PC scores (*SI Appendix* **Fig. S3D**). Models incorporating spatially explicit metrics improved cross-validation prediction of LAI and soil infiltration rate compared with models using spatial averages alone (**Figs. 3, 4, and** *SI Appendix* **Fig. S4**). The best predictive model for LAI included PC scores representing spatial averages, LMX indices, and Moran’s I, whereas that for soil infiltration rate included spatial averages and LMX indices, but not Moran’s I (*SI Appendix* **Table S4; Figs. 3 and 4**). For root biomass, the best model included spatial averages alone, but its overall explanatory performance was weak (*SI Appendix* **Fig. S4**; all-data regression: R² = 0.049, F = 1.902, P = 0.1653).

Observed versus fitted relationships using the full datasets are shown in *SI Appendix* **Fig. S4**, and statistical tests of individual predictors in the best-performing models are provided in *SI Appendix* **Table S5**.

## Discussion

Our study demonstrated that incorporating frequency-based and spatially explicit metrics of microscale nonrandomness consistently improved prediction across contrasting ecological systems compared with models based on macroscale variables or spatial averages alone. Importantly, this improvement was evaluated based on predictive performance rather than simply on model fit. The prediction targets were subsequent demographic rates in the three resource–consumer systems and macroscale functional states in the forest vegetation–soil system. Together, these results indicate that microscale nonrandomness contains predictive information beyond that captured by conventional macroscale variables and may therefore reveal ecologically relevant patterns that are obscured at broader scales.

The distributions of organisms and environmental properties are often heterogeneous at microscales and may show aggregation across physical space or among discrete biological units. In the aquatic host–parasite system, for example, variation in infection intensity represents a heterogeneous distribution of parasites among host cells, irrespective of the physical locations of individual cells. The consistent improvement in prediction associated with these microscale patterns across the four systems supports our hypothesis that nonrandom local patterns can provide information about broader ecological processes. This idea is conceptually related to spatial early warning approaches, in which nonrandom spatial patterns arising from local positive feedback can be indicators of ecological transitions, although such patterns have generally been evaluated at broader spatial scales than those considered here (Rietkerk *et al*., 2004). At finer scales, local positive feedback may similarly amplify initially small differences among biological units or locations, and these effects may subsequently propagate among units or through surrounding physical space (**Fig. 2E**). Such local amplification and propagation could underlie the temporal changes in microscale nonrandomness observed in the three resource–consumer systems (*SI Appendix* **Fig. S3A–C**) and its spatial variation in the forest vegetation–soil system (*SI Appendix* **Fig. S3D**). Importantly, our results extend the use of nonrandom spatial patterns from indicators of longer-term ecological transitions to quantitative predictors of subsequent demographic rates and macroscale functional states using microscale information.

The microscales considered in each system are biologically relevant to the processes linking local heterogeneous patterns to subsequent demographic rates or macroscale functional states. In the diatom host–chytrid parasite system, Yoneya et al. (2021) showed that infection intensity can deviate from a Poisson distribution toward an exponentially weighted Poisson distribution when existing infections promote subsequent infections, suggesting positive feedback in the spread of infection. For plant–insect systems, Winder et al. (2001) showed that predator population growth measured at 12-m intervals was positively associated with SADIE cluster indices of aphid distributions at the same spatial resolution, demonstrating that microscale insect aggregation can be linked to demographic rates of interacting species. Although this relationship differs from the cross-scale prediction of the insects’ own demographic rates in our systems, it supports the biological relevance of microscale insect distributions to demographic processes. For vegetation–soil systems, studies in semiarid ecosystems have shown close associations between microscale soil properties and vegetation patterns. Maestre & Cortina (2002) found a positive association between the spatial patterns of soil properties and vegetation based on their SADIE indices, while Maestre *et al*. (2003) showed that microscale soil properties were related to spatial variation in seedling survival. Although these studies did not examine whether the degree of microscale heterogeneity in soil properties predicts macroscale vegetation or soil functions, they support the functional relevance of microscale soil variation to vegetation–soil processes.

Unexpectedly, the frequency-based Morisita index substantially improved forecasting of subsequent demographic rates in the willow–leaf beetle and Komatsuna–aphid systems, even when spatially explicit metrics were available, and in some cases provided the best forecasts without either these spatial metrics or macroscale variables. These results have two implications. First, information contained in frequency distributions may capture an important component of microscale nonrandomness even without information on explicit spatial configuration. Although spatially explicit metrics that directly account for the locations of observations are increasingly used instead of frequency-based indices (De Freitas Alves & De Santana, 2022), our results suggest that the simpler frequency-based approach can retain substantial predictive information. Second, from a practical perspective, frequency-based indices may permit more flexible sampling designs because exact spatial relationships among sampling locations need not always be retained. This could reduce monitoring effort and broaden applicability to systems in which intensive spatial mapping is impractical. Although further investigation is needed to determine how generally this sampling advantage applies, its potential is illustrated by the komatsuna–aphid system, in which aphid population growth was best forecast by the Morisita-only model even when the reduced dataset was used to quantify predictor variables (*SI Appendix*, **Table S3**).

Our framework may provide a practical option for ecological prediction when long time series are unavailable but microscale observations can be obtained. For model construction, our framework can use short time series such as those analyzed here when supplemented by microscale observations, whereas empirical dynamic modeling (EDM) requires sufficient temporal information to reconstruct a state space and establish a library of system states (Sugihara & May, 1990; Dixon, Milicich & Sugihara, 1999). For subsequent predictions, our framework can use a single snapshot of microscale observations, whereas EDM still requires sufficient current and/or recent observations to locate the current state in the reconstructed multidimensional state space. A key remaining question concerns the sampling requirements of our framework, including the minimum number and spatial arrangement of microscale observations needed for reliable prediction and the conditions under which frequency-based metrics are sufficient or explicit spatial information is required. The spatial extent of sampling should also depend on the intended application: although the scalability of our framework to larger areas remains to be tested, large-scale extrapolation may not always be necessary where agricultural or forest management operates at relatively small local scales, as is common in Japan (Kawasaki, 2010;Morino & Horiuchi 2010). Overall, our findings highlight microscale nonrandomness as a potentially informative dimension of ecological systems, suggesting an alternative approach to the common strategy of extending monitoring over longer periods or larger spatial domains.

## Materials and methods

### Study systems and data collection

We evaluated our framework using four ecological systems differing in organisms, environments, and data structure: a laboratory diatom host–chytrid parasite system, a field willow–leaf beetle system, a greenhouse Komatsuna–aphid system, and a forest vegetation–soil system (**Fig. 1**; **see *SI Materials and Methods* for experimental and sampling details**). For the three resource–consumer systems, we repeatedly quantified population abundance of consumers and their infection or infestation prevalence together with microscale distributions of chytrid sporangia, leaf beetles, or aphids. The diatom–chytrid experiment comprised four replicate incubations monitored from day 0 to day 10. The willow–leaf beetle dataset was obtained from 48 individually potted willow plants monitored for 33 days, with population abundance and infestation prevalence summarized across plants at each census to generate a single time series, using a subset of previously published data (Yoneya *et al*., 2023). The Komatsuna– aphid experiment comprised 12 replicate units distributed among four sections within two greenhouses and was censused eight times for one month.

For the forest vegetation–soil system, we surveyed four sampling units at the Kasebaru forest site in the Shiiba Research Forest, Kyushu University. Each unit comprised three 20-m survey lines, with microscale soil surface properties measured at 0.5-m intervals. We summarized these observations over three subsections of each line corresponding to locations where macroscale functional variables were measured. The analyses presented here focused on three of these variables: leaf area index (LAI), root biomass, and soil infiltration rate. (**see *SI Materials and Methods* for the complete sampling design and measurement procedures**).

### State, rate, and heterogeneity metrics

For the three resource–consumer systems, we used population abundance (chytrid sporangia, leaf beetles, or aphids) and prevalence (the fraction of diatom host cells, willow trees, or Komatsuna plants infected or infested) as state variables. For each state variable *s*, we calculated the rate variable at time step *k* as the specific growth rate to the next time step, *τ_k_* = ln(*s_k_*_+1_/*s_k_*) /Δ*t_k_*, where Δ*t_k_* is the time interval between observations *k* and *k*+1. We set the rate to zero when both *s_k_* and *s_k_*_+1_ were zero and treated it as undefined when only one was zero. The rate was also undefined at the final observation of each time series because no subsequent observation was available or used.

To characterize microscale heterogeneity, we distinguished frequency-based heterogeneity, which describes variation in abundance among sampling units without considering their spatial arrangement, from spatially explicit heterogeneity, which incorporates the locations of those units. We quantified frequency-based heterogeneity using the Morisita index (I_δ_; Morisita (1962), which measures dispersion in counts among sampling units and is less sensitive to variation in mean abundance than the variance-to-mean ratio. We used the original I_δ_ for the diatom–chytrid system and the standardized Morisita index (Smith-Gill, 1975) for the two plant–insect systems; the rationale for using these alternative forms is provided in ***SI Materials and Methods***. We did not calculate Morisita indices for the forest vegetation–soil system because its microscale variables were binary or continuous rather than counts.

We quantified spatially explicit heterogeneity using the Li–Madden–Xu (LMX) index and Moran’s *I*. Spatial Analysis by Distance IndicEs (SADIE) characterizes spatial structure based on the minimum distance required to redistribute observed quantities among sampling locations to achieve a regular distribution (Perry, 1995). The LMX index extends this framework to characterize local patch/gap structure by quantifying the contribution of the observed value at each sampling location relative to that expected under a spatially random distribution (Li *et al*., 2012). For the two plant–insect systems, we used the overall LMX index, whereas for the forest vegetation–soil system, we averaged position-specific LMX indices within each of three subsections per survey line.

Moran’s *I* quantifies spatial autocorrelation, i.e., the similarity of observed values among spatially neighboring sampling locations (Moran, 1950). Positive values indicate that similar values tend to occur near each other, whereas negative values indicate that dissimilar values tend to be neighbors. For the two plant–insect systems, we used global Moran’s *I*, whereas for the forest vegetation–soil system, we averaged local Moran’s *I* within each of three subsections per survey line. Details of the spatial domains, neighborhood definitions, subsection averaging, and R programing implementation of both spatial metrics are provided in ***SI Materials and Methods***.

### Statistical analyses

For the diatom–chytrid, willow–leaf beetle, and Komatsuna–aphid time series, we used generalized least-squares models with a continuous autoregressive correlation structure to forecast specific growth rates *τ_k_* from time step *k* to *k*+1, using macroscale state variables and microscale heterogeneity metrics measured at time step *k* as predictors. We constructed candidate models from all combinations of the available predictors and evaluated their out-of-sample forecasting performance. For systems with replicated time series, we used leave-one-replicate-out cross-validation (LORO-CV); for the single willow–leaf beetle time series, we used rolling one-step-ahead forecasting. We primarily compared models using micro-averaged root mean squared error (RMSE), with macro-averaged RMSE and AIC reported as complementary measures (**see *SI Materials and Methods* for model fitting and cross-validation details**).

For the forest vegetation–soil system, we used a two-step analysis combining principal component analysis (PCA) and ordinary least-squares regression. We separately reduced the subsection-averaged microscale soil properties, LMX indices, and local Moran’s *I* to their first two principal components and used these six PC scores as candidate predictors of LAI, root biomass, and soil infiltration rate. We evaluated all predictor combinations using LORO-CV, with each survey line treated as a replicate. In each iteration, both PCA construction and model fitting used only the training data, and observations from the held-out line were projected onto the training PC axes before prediction (see ***SI Materials and Methods* for details**).

### Use of generative artificial intelligence

We used ChatGPT (OpenAI; GPT-5.6) to assist with manuscript editing, the generation and revision of R code, and clarification of differences among statistical methods and analytical options. We critically reviewed and tested all AI-generated or AI-revised R code and critically evaluated all AI-assisted manuscript revisions and explanations of statistical methods and analytical options. We made all final decisions regarding the manuscript, statistical methods, and analyses and take full responsibility for the final content.

## Supporting information

Supporting information including additional methods, tables and figures

## Data, Materials, and Software Availability

Original data are provided within the rendered R Notebook HTML files. All R code used for statistical analyses and visualization, except for code used to generate the SI Tables, is provided separately for each of the four study systems in the GitHub repository (https://github.com/tksmiki/microscale_nonrandomness), and also directly downloadable from the corresponding latest release (https://github.com/tksmiki/microscale_nonrandomness/releases/latest).

## Author contributions

[TM]: Conceptualization, Methodology, Formal analysis, Investigation, Project administration, Supervision, Writing – original draft, Writing – review & editing.

[KY]: Conceptualization, Methodology, Investigation, Project administration, Supervision, Writing – review & editing.

[AK]: Methodology, Investigation, Project administration, Supervision, Writing – review & editing. [HSH]: Methodology, Investigation, Writing – review & editing

[HN]: Methodology, Investigation, Writing – review & editing.

[ST]: Methodology, Investigation, Writing – review & editing.

[HM]: Investigation, Writing – review & editing.

[TK]: Conceptualization, Methodology, Investigation, Project administration, Supervision, Writing – review & editing.

[K Shirotuka]: Methodology, Project administration, Writing – review & editing.

[SK]: Methodology, Project administration, Writing – review & editing.

[K Seto]: Methodology, Resources, Writing – review & editing.

[SVdW]: Methodology, Resources, Writing – review & editing.

[MK]: Methodology, Resources, Investigation, Writing – review & editing.

## Acknowledgements and funding sources

We thank Kousuke Nagai, Mikoto Koga, Minami Okajima, Sota Nagano, and Satoru Nakamura for their assistance with fieldwork and data collection. This work was supported by Grant-in-Aid for Scientific Research (23H00538) of JSPS (to TM, KY, AK, TK, KS, SK, and MK). SVdW also was supported by the Research Council of Finland [371158].

## References

Carpenter S.R., Ludwig D. & Brock W.A. (1999). Management of Eutrophication for Lakes Subject to Potentially Irreversible Change. Ecological Applications 9, 751–771. 10.1890/1051-0761

Cheung P., Nozawa Y. & Miki T. (2021). Ecosystem engineering structures facilitate ecological resilience: A coral reef model. Ecological Research 36, 673–685. 10.1111/1440-1703.12230

Dakos V., Boulton C.A., Buxton J.E., Abrams J.F., Arellano-Nava B., Armstrong McKay D.I., et al. (2024). Tipping point detection and early warnings in climate, ecological, and human systems. Earth System Dynamics 15, 1117–1135. 10.5194/esd-15-1117-2024

Dakos V., Carpenter S.R., Van Nes E.H. & Scheffer M. (2015). Resilience indicators: prospects and limitations for early warnings of regime shifts. Philosophical Transactions of the Royal Society B: Biological Sciences 370, 20130263. 10.1098/rstb.2013.0263

Dakos V., van Nes E.H., Donangelo R., Fort H. & Scheffer M. (2010). Spatial correlation as leading indicator of catastrophic shifts. Theoretical Ecology 3, 163–174. 10.1007/s12080-009-0060-6

Das S., Shnerb N. & DeMalach N. (2026). Stochasticity and heterogeneity restrict tipping points and alternative stable states. 10.64898/2026.06.15.732263

Day C., Murphy S.T., Styles J., Taylor B., Beale T., Holland W., et al. (2024). Forecasting the population development of within-season insect crop pests in sub-Saharan Africa: the Pest Risk Information Service. Journal of Integrated Pest Management 15, 7. 10.1093/jipm/pmad026

De Freitas Alves G. & De Santana D.G. (2022). Why do traditional dispersion indices used for analysis of spatial distribution of plants tend to become obsolete? Population Ecology 64, 80–92. 10.1002/1438-390X.12105

Dixon P.A., Milicich M.J. & Sugihara G. (1999). Episodic Fluctuations in Larval Supply. Science 283, 1528–1530. 10.1126/science.283.5407.1528

Favis-Mortlock D. (1998). A self-organizing dynamic systems approach to the simulation of rill initiation and development on hillslopes. Computers & Geosciences 24, 353–372. 10.1016/S0098-3004(97)00116-7

de Groot M., Pocock M.J.O., Bonte J., Fernandez-Conradi P. & Valdés-Correcher E. (2023). Citizen Science and Monitoring Forest Pests: a Beneficial Alliance? Current Forestry Reports 9, 15–32. 10.1007/s40725-022-00176-9

Grziwotz F., Chang C.-W., Dakos V., Van Nes E.H., Schwarzländer M., Kamps O., et al. (2023). Anticipating the occurrence and type of critical transitions. Science Advances 9, eabq4558. 10.1126/sciadv.abq4558

Hastings A. (2004). Transients: the key to long-term ecological understanding? Trends in Ecology & Evolution 19, 39–45. 10.1016/j.tree.2003.09.007

Hastings A. & Wysham D.B. (2010). Regime shifts in ecological systems can occur with no warning. Ecology Letters 13, 464–472. 10.1111/j.1461-0248.2010.01439.x

Herms D.A. (2004). Using Degree-Days and Plant Phenology to Predict Pest Activity. IPM (Integrated Pest Management) of Midwest Landscapes

Hillebrand H., Donohue I., Harpole W.S., Hodapp D., Kucera M., Lewandowska A.M., et al. (2020). Thresholds for ecological responses to global change do not emerge from empirical data. Nature ecology & evolution 4, 1502–1509. 10.1038/s41559-020-1256-9

Holfeld H. (2000). Infection of the single-celled diatom Stephanodiscus alpinus by the chytrid Zygorhizidium: Parasite distribution within host population, changes in host cell size, and host– parasite size relationship. Limnology and Oceanography 45, 1440–1444. 10.4319/lo.2000.45.6.1440

Hurlbert S.H. (1990). Spatial Distribution of the Montane Unicorn. Oikos 58, 257–271. 10.2307/3545216

Jiménez D., Delerce S., Dorado H., Cock J., Muñoz L.A., Agamez A., et al. (2019). A scalable scheme to implement data-driven agriculture for small-scale farmers. Global Food Security 23, 256–266. 10.1016/j.gfs.2019.08.004

Kaneko S., Shirotsuka K. & Shibao M. (2017). Forecast of peak dates of adult emergence of Helicoverpa armigera in Osaka Prefecture by using a simulation program based on the total effective temperature provided by JPP-NET. Annual Report of the Kanwai Plant Protection Society 59, 105–108. 10.4165/kapps.59.105

Kawasaki K. (2010). The costs and benefits of land fragmentation of rice farms in Japan. Australian Journal of Agricultural and Resource Economics 54, 509–526. 10.1111/j.1467-8489.2010.00509.x

Kéfi S., Guttal V., Brock W.A., Carpenter S.R., Ellison A.M., Livina V.N., et al. (2014). Early Warning Signals of Ecological Transitions: Methods for Spatial Patterns. PLoS ONE 9, e92097. 10.1371/journal.pone.0092097

Kidd D. & Amarasekare P. (2012). The role of transient dynamics in biological pest control: insights from a host–parasitoid community. Journal of Animal Ecology 81, 47–57. 10.1111/j.1365-2656.2011.01881.x

Kuehn C. (2011). A mathematical framework for critical transitions: bifurcations, fast-slow systems and stochastic dynamics. Physica D: Nonlinear Phenomena 240, 1020–1035. 10.1016/j.physd.2011.02.012

Legendre P. & Fortin M.J. (1989). Spatial pattern and ecological analysis. Vegetatio 80, 107–138. 10.1007/BF00048036

Li B., Madden L.V. & Xu X. (2012). Spatial analysis by distance indices: an alternative local clustering index for studying spatial patterns: *A new SADIE clustering index*. Methods in Ecology and Evolution 3, 368–377. 10.1111/j.2041-210X.2011.00165.x

Lloyd-Smith J.O., Schreiber S.J., Kopp P.E. & Getz W.M. (2005). Superspreading and the effect of individual variation on disease emergence. Nature 438, 355–359. 10.1038/nature04153

Maestre F.T. & Cortina J. (2002). Spatial patterns of surface soil properties and vegetation in a Mediterranean semi-arid steppe. Plant and Soil 241, 279–291. 10.1023/A:1016172308462

Maestre F.T., Cortina J., Bautista S., Bellot J. & Vallejo R. (2003). Small-scale Environmental Heterogeneity and Spatiotemporal Dynamics of Seedling Establishment in a Semiarid Degraded Ecosystem. Ecosystems 6, 630–643. 10.1007/s10021-002-0222-5

Miki T., Ushio M., Fukui S. & Kondoh M. (2010). Functional diversity of microbial decomposers facilitates plant coexistence in a plant–microbe–soil feedback model. Proceedings of the National Academy of Sciences 107, 14251–14256. 10.1073/pnas.0914281107

Moran P.A.P. (1950). Notes on Continuous Stochastic Phenomena. Biometrika 37, 17–23. 10.2307/2332142

Morino M., Horiuchi S. (2010). Effects of Location on the Management of Small Private Forests. Studies in Regional Science, 40, 143–155

Morisita M. (1962). I_delta-INDEX, A MEASURE OF DISPERSION OF INDIVIDUALS. Research in Population Ecology, 1–7

Mumby P.J., Hastings A. & Edwards H.J. (2007). Thresholds and the resilience of Caribbean coral reefs. Nature 450, 98–101. 10.1038/nature06252

O’Brien D.A., Deb S., Gal G., Thackeray S.J., Dutta P.S., Matsuzaki S.S., et al. (2023). Early warning signals have limited applicability to empirical lake data. Nature Communications 14, 7942. 10.1038/s41467-023-43744-8

Pauly D. & Zeller D. (2016). Catch reconstructions reveal that global marine fisheries catches are higher than reported and declining. Nature Communications 7, 10244. 10.1038/ncomms10244

Perry J.N. (1998). Measures of Spatial Pattern for Counts. Ecology 79, 1008–1017. 10.2307/176596

Perry J.N. (1995). Spatial Analysis by Distance Indices. Journal of Animal Ecology 64, 303–314. 10.2307/5892

Rietkerk M., Dekker S.C., De Ruiter P.C. & Van De Koppel J. (2004). Self-Organized Patchiness and Catastrophic Shifts in Ecosystems. Science 305, 1926–1929. 10.1126/science.1101867

Scheffer M., Bascompte J., Brock W.A., Brovkin V., Carpenter S.R., Dakos V., et al. (2009). Early-warning signals for critical transitions. Nature 461, 53–59. 10.1038/nature08227

Smith-Gill S.J. (1975). Cytophysiological basis of disruptive pigmentary patterns in the leopard frogRana pipiens. II. Wild type and mutant cell-specific patterns,. Journal of Morphology 146, 35–54. 10.1002/jmor.1051460103

Stein R.A. (2011). Super-spreaders in infectious diseases. International Journal of Infectious Diseases 15, e510–e513. 10.1016/j.ijid.2010.06.020

Su H., Wang R., Feng Y., Li Y., Li Y., Chen J., et al. (2021). Long-term empirical evidence, early warning signals and multiple drivers of regime shifts in a lake ecosystem. Journal of Ecology 109, 3182–3194. 10.1111/1365-2745.13544

Sugihara G. & May R.M. (1990). Nonlinear forecasting as a way of distinguishing chaos from measurement error in time series. Nature 344, 734–741. 10.1038/344734a0

Turechek W.W. & Madden L.V. (1999). Spatial Pattern Analysis of Strawberry Leaf Blight in Perennial Production Systems. Phytopathology® 89, 421–433. 10.1094/PHYTO.1999.89.5.421

Weinans E., Quax R., Van Nes E.H. & Leemput I.A.V.D. (2021). Evaluating the performance of multivariate indicators of resilience loss. Scientific Reports 11, 9148. 10.1038/s41598-021-87839-y

Winder L., Alexander C., Griffiths G., Holland J., Wooley C. & Perry J. (2019). Twenty years and counting with SADIE: Spatial Analysis by Distance Indices software and review of its adoption and use. Rethinking Ecology 4, 1–16. 10.3897/rethinkingecology.4.30890

Winder L., Alexander C.J., Holland J.M., Woolley C. & Perry J.N. (2001). Modelling the dynamic spatio-temporal response of predators to transient prey patches in the field. Ecology Letters 4, 568–576. 10.1046/j.1461-0248.2001.00269.x

Yoneya K., Miki T. & Takabayashi J. (2023). Initial herbivory and exposure to herbivory-induced volatiles enhance arthropod species richness by diversifying community assemblages. Frontiers in Ecology and Evolution 10, 1031664. 10.3389/fevo.2022.1031664

Yoneya K., Miki T., Van den Wyngaert S., Grossart H. & Kagami M. (2021). Non-random patterns of chytrid infections on phytoplankton host cells: mathematical and chemical ecology approaches. Aquatic Microbial Ecology 87, 1–15. 10.3354/ame01966

Żychaluk K., Bruno J.F., Clancy D., McClanahan T.R. & Spencer M. (2012). Data-driven models for regional coral-reef dynamics: Data-driven coral-reef models. Ecology Letters 15, 151–158. 10.1111/j.1461-0248.2011.01720.x

Morino M. & Horiuchi S. (2010). Effects of location on the management of small private forests. Studies in Regional Science 40, 143–155. 10.2457/srs.40.143

