## Supporting information including additional methods, tables and figures for "Microscale nonrandomness revisited: An overlooked predictor of ecological change"


#### This PDF file includes:

SI Materials and Methods

Tables S1-S5

Figures S1-S5

SI References

### SI Materials and Methods

#### Study systems and data collection

##### Diatom host–chytrid parasite system

Fungal parasites such as chytrids are important components of freshwater food webs and can influence phytoplankton population dynamics and diversity as well as the transfer of matter to higher trophic levels (Kagami, Helmsing & van Donk, 2011; Miki, Takimoto & Kagami, 2011; Frenken *et al.*, 2017). Chytrid infections are also a major concern in large-scale algal cultivation for biofuel production, where preventing their spread is important for maintaining stable cultures (Smith *et al.*, 2015; Widin *et al.*, 2022). Thus, monitoring and forecasting chytrid dynamics are important in both natural and artificial systems. We conducted four replicate incubations, each containing 30 mL of WC medium (Guillard & Lorenzen, 1972; Andersen *et al.*, 2005) in a 50-mL VIOLAMO Centrifuge Tube II (VIO-50BN; AS ONE Corporation, Japan). We last subcultured the diatom inoculum (*Ulnaria sp.* SYN2) and the diatom–chytrid (*Zygophlyctis planktonicum* SYN-CHY1) culture (Seto *et al.*, 2020) used as the chytrid source on 14 May 2026, 11 days before the experiment. We initiated the experiment on 25 May 2026 (day 0). Based on the diatom density measured on day 0 before inoculation (15,796 cells mL$^{-1}$), we added 100 µL of the diatom inoculum to each incubation to obtain an initial density of approximately 50 cells mL$^{-1}$ (the calculated volume was 94.96 µL, which we rounded to 100 µL for ease of pipetting). On day 0, the diatom–chytrid culture used as the chytrid source had a host density of 1,320 cells mL$^{-1}$ and an infection prevalence of 0.430. To minimize potential damage to chytrid zoospores, we allowed the culture to stand without centrifugation so that host cells settled naturally toward the bottom. We then collected 100 µL of the upper liquid fraction, which was expected to contain chytrid zoospores while minimizing the transfer of host cells, and added it to each incubation as the chytrid inoculum.

Beginning on day 0 and continuing through day 10, we subsampled 1.4 mL from each incubation and preserved each sub-sample with 14 µL of Lugol's solution (pH 7). Before microscopic examination, we added 20 µL of 0.1 mol L$^{-1}$ sodium thiosulfate to remove the brown coloration caused by Lugol's solution and subsequently stained the samples with BD Calcofluor White Reagent Dropper (CFW; Becton Dickinson, Maryland, USA). We examined host cells and chytrid infections using a CX43 microscope (Evident) equipped with a UV fluorescence unit (LED-FL3/UV, 355 nm; AdvanVision). For each 1.4-mL subsample, we examined all host cells in 1.0 mL of the subsample and counted all chytrid sporangia attached to each host cell, including immature, mature, and empty sporangia, without distinguishing among these categories. We used the number of sporangia per host cell as a measure of infection intensity. We excluded empty host cells, regardless of whether chytrid sporangia were attached, because we considered these cells remnants of previous infections. Although some sporangia attached to empty host cells may remain infective, we excluded them because it was difficult to distinguish empty from non-empty sporangia. We also excluded host cells surrounded by extensive CFW-stained fungal structures following leakage of cellular contents because we considered these structures to represent advanced remnants of previous infections and could not reliably distinguish and count individual sporangia.

##### Willow–leaf beetle system

Willows harbor diverse arthropod communities and provide a basis for their interactions in riparian ecosystems (Utsumi, Nakamura & Ohgushi, 2009). Among the many herbivores associated with willows, leaf beetle herbivory induces chemical and physical responses in host plants that can influence subsequent community assembly and, consequently, biodiversity (Yoneya, Miki & Katayama, 2023a; Yoneya, Miki & Takabayashi, 2023b). Thus, monitoring and forecasting leaf beetle population dynamics may help anticipate their broader consequences for riparian arthropod communities. We reused a subset of data from a published study (Yoneya *et al.*, 2023b) that investigated the initial colonization and population dynamics of the leaf beetle *Plagiodera versicolora*, as well as the initial assembly of arthropod communities on the willow *Salix eriocarpa*. After three pretreatments (undamaged, damaged by *P. versicolora*, and exposed to volatiles emitted by damaged conspecifics), we transferred 48 individually potted willow plants (16 per pretreatment) to the experimental garden at the Center for Ecological Research, Kyoto University, at 06:00 on day 1 (10 June 2008). We arranged the potted plants in two parallel lines, with 1 m between adjacent plants within each line and 2 m between the two lines.

We used census data on *P. versicolora* abundance collected at varying intervals over the course of the experiment. On day 1, we censused the plants hourly from 07:00 to 12:00 (six censuses) and then every 2 h from 14:00 to 18:00 (three censuses). We conducted another census at 07:00 on day 2 and subsequently at 09:00 and 16:00 each day from day 2 to day 7. Thereafter, we conducted censuses at 09:00 and 16:00 on days 10 and 32, at 09:00 on day 33, at 09:00 and 16:00 on day 60, and at 09:00 on day 61. For the time-series analysis, we excluded the final three abundance observations (two on day 60 and one on day 61) because of the substantially longer interval between days 33 and 60 than the preceding census intervals. In the present study, we analyzed only the abundance of adult *P. versicolora*.

**Komatsuna–aphid system**

The peach green aphid, *Myzus persicae*, is a cosmopolitan pest of diverse crops, including Brassicaceae and Solanaceae, and rapidly evolves resistance to insecticides (Bass *et al.*, 2014). Monitoring and forecasting its population dynamics can therefore facilitate timely pest management while reducing reliance on insecticides and improving the effective use of biological control agents, such as ladybugs (Beltrà *et al.*, 2018).We conducted the experiment in two greenhouses at the Research Institute of Environment, Agriculture, Forestry and Fisheries, Osaka, located sufficiently far apart to minimize interactions between them. We partitioned each greenhouse into northern and southern sections using a plastic curtain positioned at the center. Each section consisted of four north–south-oriented ridges spaced 110 cm apart. We left the easternmost ridge unplanted and established two parallel planting lines, separated by 25 cm, on each of the remaining three ridges.

We prepared Komatsuna (*Brassica rapa* var. *perviridis* cv. Hitomi) seedlings 5 days before transplantation. To establish initial aphid colonies before the experiment began, we inoculated each of four seedlings with three individuals of the aphid (*M. persicae*) in an incubator (25°C, 16L8D) and kept the remaining seedlings aphid-free. On 1 April 2024, we transplanted all seedlings at 10-cm intervals, with 65 seedlings in each planting line. We numbered the six planting lines consecutively from west to east (Lines 1–6) and the seedlings within each line from north to south (Individuals 1–65). We placed the four aphid-inoculated seedlings at Individuals 32 and 65 of Line 1 and Individuals 1 and 65 of Line 3. Although we kept both ends of each greenhouse closed throughout the experiment, the greenhouse structure allowed natural immigration of aphids from the surrounding environment. To minimize edge effects, we included only Individuals 23–42 of each line in subsequent censuses. We conducted censuses on 4, 8, 12,

15, 19, 22, 26, and 30 April 2024 between 09:00 and 14:00.

For visualization of aphid spatial distributions (**Fig. 2D and Fig. S2**), we rearranged the four sections horizontally rather than preserving their actual 2 × 2 spatial arrangement. From left to right, the displayed sections corresponded to Section 1 (northern section of the first greenhouse), Section 2 (southern section of the first greenhouse), Section 3 (northern section of the second greenhouse), and Section 4 (southern section of the second greenhouse). Within each section, the x- and y-coordinates represented positions relative to the upper-left corner of that section and preserved the relative spatial arrangement of plants. We set the horizontal distances between sections arbitrarily for visualization; thus, these distances did not represent the actual distances between sections or greenhouses. We calculated all microscale spatial indices separately within each section, so these artificial between-section distances did not affect the spatial analyses.

**Forest vegetation–soil system**

Many forests in Japan have experienced vegetation degradation associated with sika deer overpopulation, leading to soil erosion, suppressed tree growth rate, and reduced carbon stocks and potentially reinforcing further vegetation degradation (Harada, Ang Meng Ann & Suzuki, 2020; Abe *et al.*, 2024a; Abe, Kume & Katayama, 2024b). Because vegetation degradation and soil erosion can emerge locally, simple methods for evaluating current vegetation and soil conditions could facilitate low-cost assessment of forest degradation from readily observable microscale properties. We conducted field surveys in three forest plots in the Shiiba Research Forest, Kyushu University; however, because of substantial differences in environmental conditions among the plots, particularly in vegetation composition and soil parent material, we used only data collected from Plot 1 in the present study. We established Plot 1 at the Kasebaru forest site in Miyazaki, Japan (32.38°N, 131.10°E). This site is part of the Monitoring Sites 1000 Project, a nationwide long-term ecological monitoring program initiated by the Ministry of the Environment of Japan (Ishihara *et al.*, 2010). The site comprises a temperate mixed forest dominated by deciduous broad-leaved species, including *Acer sieboldianum* and *Carpinus laxiflora*, and evergreen conifers, such as *Tsuga sieboldii*. The plot consisted of four sampling units (Units 1–4), each containing three parallel survey lines (Lines 1–3). We separated adjacent units by 10 m and adjacent survey lines within each unit by 2 m. Each survey line extended 20 m downslope from a predefined starting point at the upper edge of the slope. We established census points at 0.5-m intervals along each line, yielding 41 census points (S01–S41).

We measured microscale soil surface properties at every census point (**Fig. S5**) and macroscale functional variables at three census points (S07, S21, and S35) along each survey line (**Fig. 1D**). The microscale soil surface properties included the presence or absence of soil pillars and exposed roots, litter layer thickness (cm), soil hardness (mm), and percentage cover (%) of litter, vegetation (e.g., tree seedlings and mosses), and stones, and total cover (%), calculated as the sum of litter, vegetation, and stone cover. We measured soil hardness directly at each census point using a Yamanaka-type soil hardness tester (DIK-5553, Daiki Rika Kogyo Co., Saitama, Japan), whereas we assessed the remaining microscale variables within a 20 × 20 cm quadrat placed adjacent to each census point. We conducted the survey of microscale soil surface properties on 8 September 2024.

The macroscale functional variables included leaf area index (LAI; $m^2$ $m^{-2}$), mass of soil components (g 100 $cm^{-3}$; minerals, roots, stones, and other components), soil bulk density (g $cm^{-3}$), soil infiltration rate (mm $h^{-1}$), and eroded soil depth (cm). We measured LAI, mass of soil components and bulk density, and infiltration rate on 28 November 2024, 27 November 2024, and 15 December 2024, respectively. We quantified LAI from fisheye images obtained using a 360° camera (Insta360 X4, Insta360, Shenzhen, China) and quantified soil composition from 100-$cm^3$ soil samples collected at each location. We quantified eroded soil depth as the change in soil surface elevation between 28 November 2024 and 11 June 2025 using the pin method. We also evaluated microbial multifunctionality as a macroscale functional variable using the BIOLOG EcoPlate assay (Miki *et al.*, 2018); however, we did not include these data in the present study.

#### Metric calculations

##### State and rate variables

We defined the state and rate variables as described in the main-text Methods. Specifically, except for the forest vegetation–soil system, population abundance referred to chytrid sporangia, adult leaf beetles, or aphids, and prevalence referred to the fraction of diatom host cells, willow trees, or Komatsuna plants infected or infested, respectively. For a state variable $s$, we calculated

$$r_k = \frac{\ln(s_{k+1}/s_k)}{\Delta t_k},$$

where $\Delta t_k$ is the time interval between observations at time step i and $k$ + 1.
We set $r_k$ = 0 when both $s_k$ and $s_{k+1}$ were zero, treated $r_k$ as undefined when only one was zero, and did not calculate it for the final observation because a subsequent observation was unavailable or not used.

##### Frequency-based heterogeneity metrics

As described in the main text, we used the Morisita index ($I_\delta$; Morisita, 1962) to quantify frequency-based heterogeneity in count data. The index is defined as

$$I_\delta = q\frac{\sum_{i=1}^{q} x_i\,(x_i - 1)}{N(N-1)}, \qquad N = \sum_{i=1}^{q} x_i,$$

where $x_i$ is the number of individuals in the $i$th sampling unit ($i = 1,2,\dots,q$), $q$ is the number of sampling units, and $N$ is the total number of individuals.

In the diatom–chytrid system, each host cell represented a sampling unit; thus, the number of sampling units varied over time with host abundance. Because the relative sensitivity of the original and standardized Morisita indices to the number of sampling units remains debated, we initially considered both. However, in the standardized Morisita index (Smith-Gill, 1975), the denominator of the term used to standardize deviations from the threshold value of 0.5 increases with the number of sampling units. With the large and variable numbers of host cells in our data, this standardization compressed values toward 0.5 even when $I_\delta$ was substantially greater than 1, potentially reducing its ability to capture variation useful for prediction. We therefore used the original $I_\delta$ for the diatom–chytrid system and the standardized Morisita index for the two plant–insect systems, in which the number of sampling units (plant individuals) remained constant over time. We calculated both metrics using the *dispindmorisita* function in the R package *vegan*. We did not calculate Morisita indices for the forest vegetation–soil system because its microscale variables were binary or continuous rather than counts.

**Spatially explicit heterogeneity metrics**

For both SADIE/LMX and Moran's *I*, we calculated the spatial metrics across all sampled plants for the willow–leaf beetle system, separately for each of the 12 replicate units (three units in each of four sections) for the Komatsuna–aphid system, and separately for each of the four sampling units (each comprising three survey lines) for the forest vegetation–soil system.

As described in the main text, SADIE evaluates spatial structure based on the minimum distance required to redistribute observed quantities among sampling locations to achieve a regular distribution (Perry, 1995), and the LMX index characterizes local patch/gap structure relative to that expected under a spatially random distribution (Li *et al.*, 2012). We calculated both the overall and position-specific LMX indices using the *sadie* function in the R package *epiphy*. For the willow–leaf beetle and Komatsuna–aphid systems, we used the overall LMX index at each time point. For the forest vegetation–soil system, we obtained three representative values for each survey line by averaging the local LMX indices over S01–S14, S15–S28, and S29–S41, respectively.

For Moran's *I* (Moran, 1950), we defined sampling locations as neighbors when they were within $3\sqrt{2}$m, $20\sqrt{2}$ cm, and $2\sqrt{2}$ m of each other for the willow–leaf beetle, Komatsuna–aphid, and forest vegetation–soil systems, respectively. We calculated global Moran's *I* using the *moran.test* function and position-specific local Moran's *I* using the *localmoran* function in the R package *spdep*. For the willow–leaf beetle and Komatsuna–aphid systems, we used global Moran's *I* at each time point. For the forest vegetation–soil system, we first calculated local Moran's *I* at each sampling point within each sampling unit and then obtained three representative values for each survey line by averaging the local indices over S01–S14, S15–S28, and S29–S41, respectively.

#### Statistical analyses

**Forecasting in time-series systems**

For the three systems with time-series data (diatom–chytrid, willow–leaf beetle, and Komatsuna–aphid), we fitted generalized least-squares (GLS) models with a continuous autoregressive correlation structure (*corCAR1*) to account for temporal autocorrelation with unequal observation intervals, using the *gls* function in the R package *nlme*. We used maximum likelihood (ML) when fitting models to training data and restricted maximum likelihood (REML) when fitting models to the full dataset. For each response variable, we constructed a candidate model set comprising all combinations of macroscale and microscale variables measured at time step *k* as predictors of the specific growth rate from *k* to *k* + 1 The macroscale predictors were diatom abundance, chytrid sporangia abundance, and infection prevalence for the diatom–chytrid system; adult leaf beetle abundance per plant and infestation prevalence for the willow–leaf beetle system; and aphid abundance and infestation prevalence for the Komatsuna–aphid system. The microscale predictors were the Morisita index $I_\delta$ for the diatom–chytrid system and the standardized Morisita index, global Moran's *I*, and overall LMX index for the two plant–insect systems.

We evaluated forecasting performance using leave-one-replicate-out cross-validation (LORO-CV) for the diatom–chytrid and Komatsuna–aphid systems, for which replicate time series were available. Each iteration used three replicates for training and one for testing in the diatom–chytrid system, and 11 for training and one for testing in the Komatsuna–aphid system. In the latter system, very low aphid abundance (e.g., zero or one individual in a section) sometimes prevented the calculation of specific growth rates or nonrandomness metrics, resulting in shorter usable time series than the original census series. We therefore excluded Unit 3 of Section 2 from testing because its usable time series contained fewer than three observations. For the single time series of the willow–leaf beetle system, we instead evaluated forecasting performance using rolling one-step-ahead forecasts over the final six time points. We selected models primarily according to out-of-sample forecasting performance rather than AIC because our primary interest was generalization to new observations; nevertheless, we also calculated and reported AIC for each model fitted to the training data. We primarily quantified forecasting performance using micro-averaged root mean squared error (RMSE), which weights individual observations equally across replicates, and additionally reported macro-averaged RMSE, which weights replicates equally, to assess robustness across replicates.

##### Prediction in the forest vegetation–soil system

For the forest vegetation–soil system, we first reduced the dimensionality of the microscale variables using principal component analysis (PCA; **Fig. S3D**). We conducted separate PCAs for three groups of variables: (1) the subsection-averaged values of the eight microscale soil properties (**Fig. S5**), (2) the subsection-averaged LMX indices, and (3) the subsection-averaged local Moran's *I*. We retained the first two PC scores from each PCA, yielding six candidate predictors: $PC1_{mean}$, $PC2_{mean}$, $PC1_{LMX}$, $PC2_{LMX}$, $PC1_{Moran}$, and $PC2_{Moran}$.

We then fitted ordinary least-squares (OLS) multiple regression models to predict each of three macroscale functional variables (LAI, root biomass, and soil infiltration rate) and constructed a candidate model set comprising all combinations of the six PC predictors. We evaluated predictive performance using the same LORO-CV framework as for the time-series systems, treating each survey line as a replicate. In each iteration, we conducted the three PCAs using only the 33 observations from the 11 training lines and projected the three observations from the held-out line onto the resulting PC axes. We fitted the OLS models using the PC scores obtained from the training-data PCA and generated predictions for the held-out observations using their projected PC scores as predictors. We also calculated and reported AIC for each model fitted to the training data.

#### Tables S1-S5

Because the model-comparison tables are extensive, we provide them as interactive HTML files. Please download the files using the links below and open them in a web browser (e.g., Google Chrome or Microsoft Edge).

##### Table S1 Model comparison for the diatom host–chytrid parasite system

Model comparisons based on forecasting performance are provided for population growth rate and prevalence growth rate. The tables also identify the predictors included in the baseline models, best-performing models, and alternative models incorporating microscale nonrandomness metrics.

**[DOWNLOAD]**

https://github.com/tksmiki/microscale_nonrandomness/releases/download/v0.1/TableS01_micro.html

**Table S2 Model comparison for the willow–leaf beetle system**

Model comparisons based on forecasting performance are provided for population growth rate and prevalence growth rate. The tables also identify the predictors included in the baseline models, best-performing models, and alternative models incorporating microscale nonrandomness metrics.

**[DOWNLOAD]**

https://github.com/tksmiki/microscale_nonrandomness/releases/download/v0.1/TableS02_micro.html

**Table S3 Model comparison for the Komatsuna–aphid system**

Model comparisons based on forecasting performance are provided for population growth rate and prevalence growth rate. The tables also identify the predictors included in the baseline models, best-performing models, and alternative models incorporating microscale nonrandomness metrics.

**[DOWNLOAD]**

https://github.com/tksmiki/microscale_nonrandomness/releases/download/v0.1/TableS03_micro.html

**Table S4 Model comparison for the forest vegetation–soil system**

Model comparisons based on predictive performance are provided for three macroscale functional variables: LAI, root biomass, and soil infiltration rate. The tables also identify the predictors included in the baseline models, best-performing models, and alternative models incorporating microscale nonrandomness metrics.

**[DOWNLOAD]**

https://github.com/tksmiki/microscale_nonrandomness/releases/download/v0.1/TableS04_micro.html

**Table S5 Summary of the best-performing models across systems**

Statistics for the best-performing models among all candidate combinations of predictors are summarized for each system.

**[DOWNLOAD]**

https://github.com/tksmiki/microscale_nonrandomness/releases/download/v0.1/TableS05_micro.html

#### Figures S1-S5

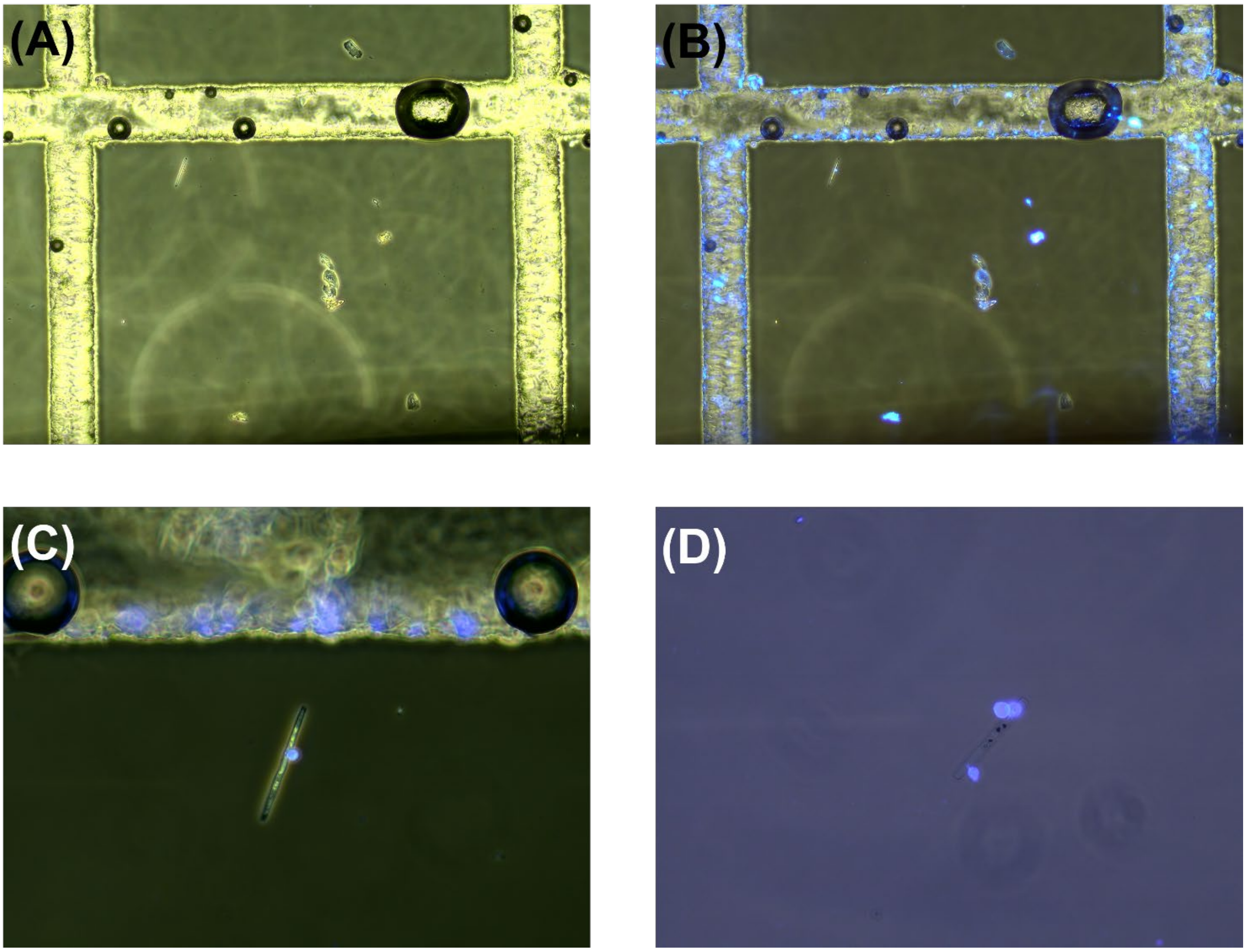


**Figure S1. Examples of different infection intensities of chytrids on individual diatom cells.** (A) Phase-contrast micrograph taken using a 10× objective lens and a plankton counting plate (Rigo-sha, model 5608-C). A single chytrid sporangium attached to the diatom cell near the upper left corner of the grid would be recorded as an infection intensity of 1. (B, C) Fluorescence micrographs of the same cell shown in (A), taken using 10× (B) and 40× (C) objective lenses. Images in (A–C) were obtained on day 3 from replicate WC1 specifically for illustration, after all observations used in the analyses had been completed. Therefore, data from this cell were not included in the analyses. (D) Fluorescence micrograph taken using a 40× objective lens. The diatom cell was recorded as an infection intensity of 3, corresponding to three attached chytrid sporangia. This image was obtained on day 3 from replicate WC2, and the cell was included in the analyses. The grid spacing in (A) and (B) is 1,000 µm, measured between the centerlines of adjacent grid-separating bands, providing a size reference. None of the images were cropped, and their relative dimensions were preserved. The original photographs (IMG_000275.jpg, IMG_000276.jpg, IMG_000277.jpg, and IMG_000137.jpg, corresponding to panels A–D, respectively) are available at https://github.com/tksmiki/microscale_nonrandomness.

**[DOWNLOAD]**

https://github.com/tksmiki/microscale_nonrandomness/releases/download/v0.1/figureS02_micro.html

**Figure S2. Spatiotemporal distributions in two plant–insect systems.** Spatiotemporal distributions of organisms and position-specific LMX indices are shown as animated GIFs; therefore, the figure is provided in HTML format. (A) Spatial distribution of adult leaf beetles among willow trees and its temporal dynamics. Each grid cell represents a single willow tree. (B) Spatial distribution of position-specific LMX indices and its temporal dynamics, shown as an example from the willow–leaf beetle system. (C) Spatial distribution of aphids on Komatsuna and its temporal dynamics.

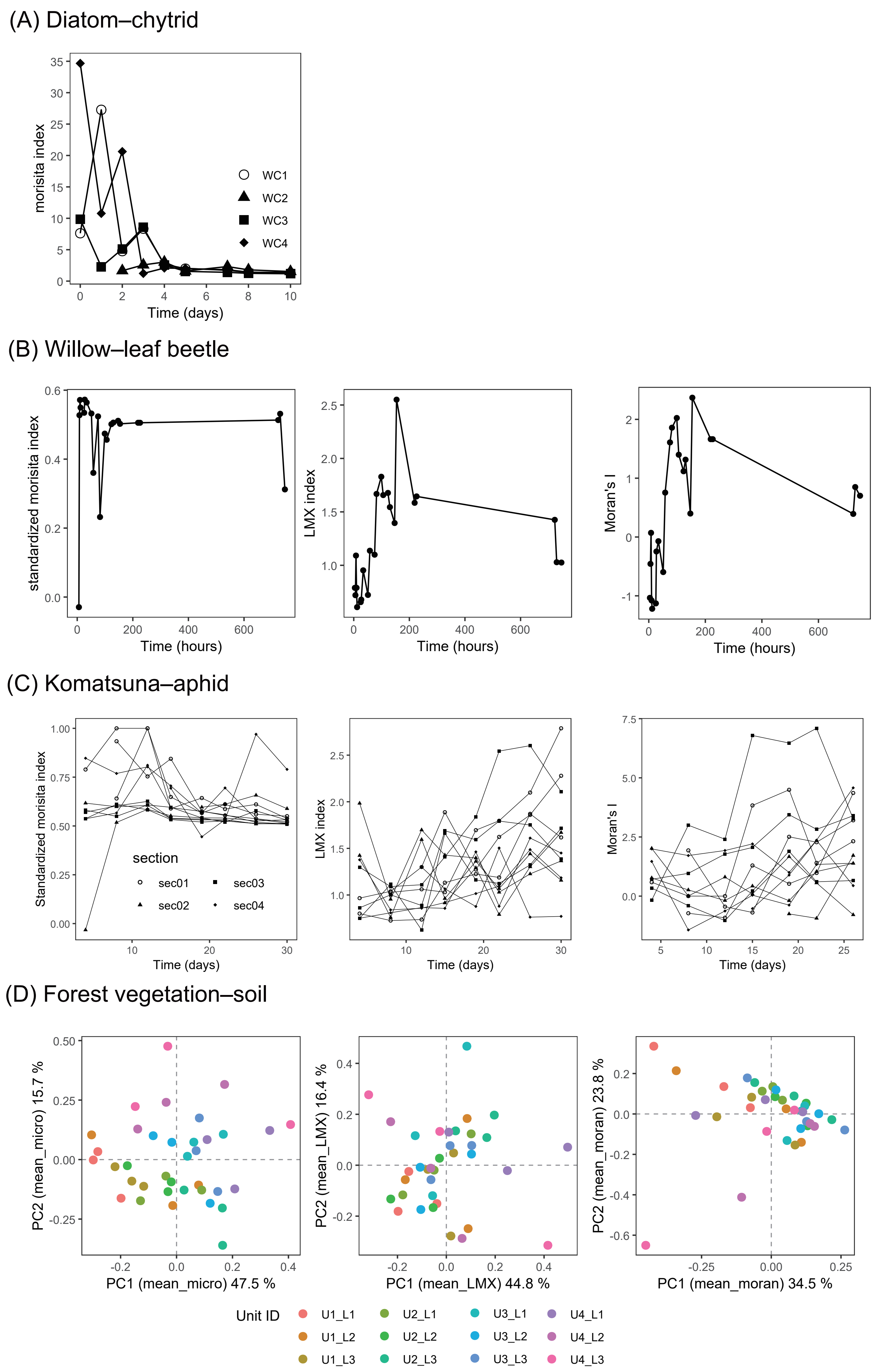

(A) Diatom–chytrid
morisita index
Time (days)
WC1
WC2
WC3
WC4
(B) Willow–leaf beetle
standardized morisita index
LMX index
Moran's I
Time (hours)
(C) Komatsuna–aphid
Standardized morisita index
LMX index
Moran's I
Time (days)
section
sec01
sec02
sec03
sec04
(D) Forest vegetation–soil
PC2 (mean_micro) 15.7 %
PC1 (mean_micro) 47.5 %
PC2 (mean_LMX) 16.4 %
PC1 (mean_LMX) 44.8 %
PC2 (mean_moran) 23.8 %
PC1 (mean_moran) 34.5 %
Unit ID
U1_L1
U1_L2
U1_L3
U2_L1
U2_L2
U2_L3
U3_L1
U3_L2
U3_L3
U4_L1
U4_L2
U4_L3

**Figure S3. Temporal dynamics and spatial distribution of microscale metrics.** (A) Temporal dynamics of Morisita ($I_\delta$) index in the diatom–chytrid system. Temporal dynamics of the standardized Morisita index, LMX index, and Moran's *I* in (B) the willow–leaf beetle system and (C) the Komatsuna–aphid system. (D) PCA scores for soil properties, obtained separately from the spatial averages of the eight soil variables, their corresponding LMX indices, and their corresponding Moran's *I* values. For visualization in this panel, each PCA was performed using all 36 observations. In the cross-validation analyses, however, PCA was performed separately for each training set (33 observations), and the held-out observations were projected onto the resulting PC axes (see **SI Materials and Methods**).

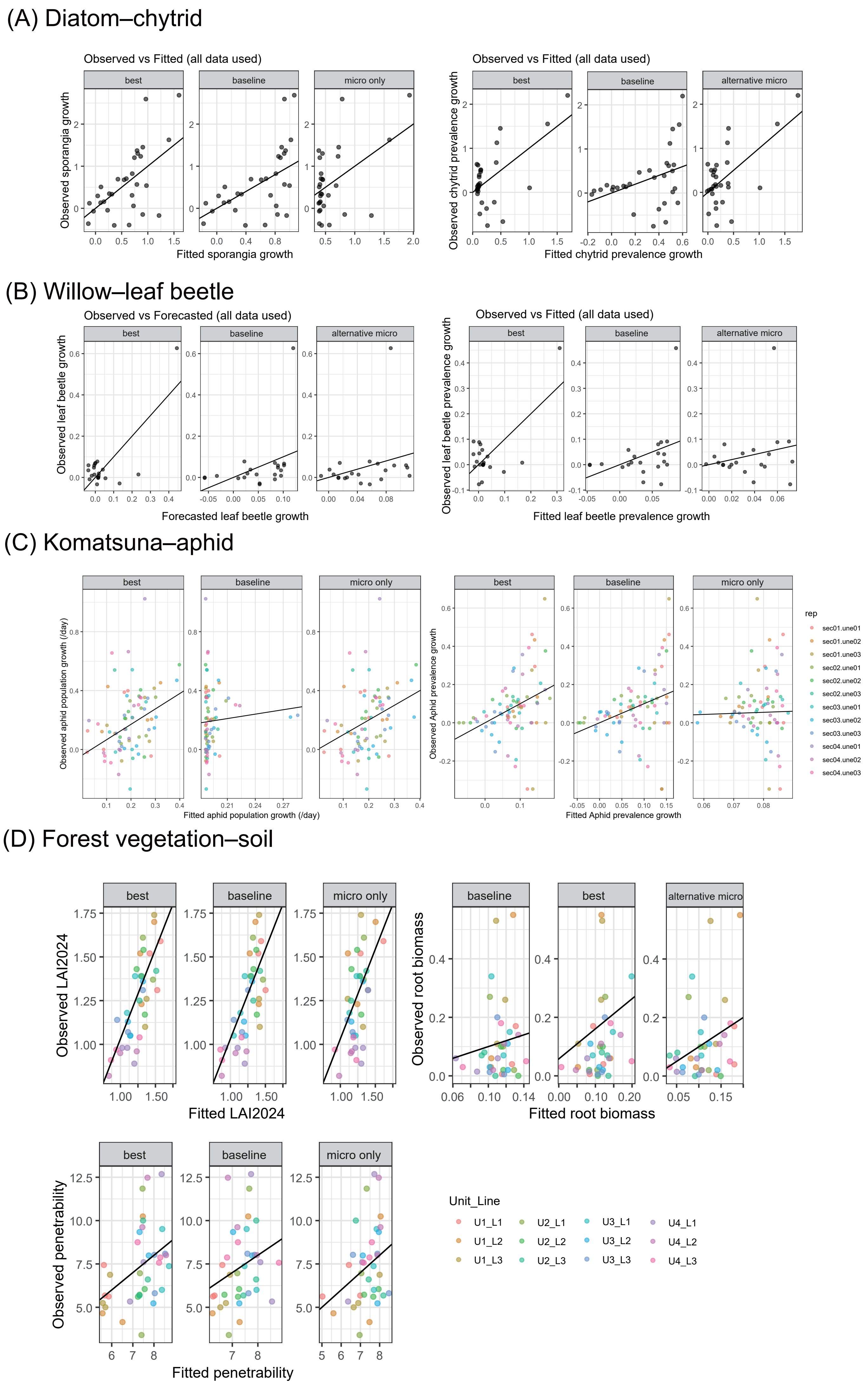

(A) Diatom–chytrid
Observed vs Fitted (all data used)
best
baseline
micro only
Observed sporangia growth
Fitted sporangia growth
Observed vs Fitted (all data used)
best
baseline
alternative micro
Observed chytrid prevalence growth
Fitted chytrid prevalence growth
(B) Willow–leaf beetle
Observed vs Forecasted (all data used)
best
baseline
alternative micro
Observed leaf beetle growth
Forecasted leaf beetle growth
Observed vs Fitted (all data used)
best
baseline
alternative micro
Observed leaf beetle prevalence growth
Fitted leaf beetle prevalence growth
(C) Komatsuna–aphid
best
baseline
micro only
Observed aphid population growth (/day)
Fitted aphid population growth (/day)
best
baseline
micro only
Observed Aphid prevalence growth
Fitted Aphid prevalence growth
rep
sec01.une01
sec01.une02
sec01.une03
sec02.une01
sec02.une02
sec02.une03
sec03.une01
sec03.une02
sec03.une03
sec04.une01
sec04.une02
sec04.une03
(D) Forest vegetation–soil
best
baseline
micro only
Observed LAI2024
Fitted LAI2024
baseline
best
alternative micro
Observed root biomass
Fitted root biomass
best
baseline
micro only
Observed penetrability
Fitted penetrability
Unit_Line
U1_L1
U1_L2
U1_L3
U2_L1
U2_L2
U2_L3
U3_L1
U3_L2
U3_L3
U4_L1
U4_L2
U4_L3

**Figure S4. Comparisons between observed and fitted target variables using all observations.** For each target variable, the "baseline" panel shows results from the best-performing model among those without microscale nonrandomness metrics. The "best" panel shows the best-performing model among those incorporating microscale nonrandomness metrics; these models were also the overall best-performing models for all target variables except root biomass, for which the baseline model performed best overall. "micro only" and "alternative micro " show alternative models incorporating microscale nonrandomness metrics. Model selection was based on out-of-sample forecasting or predictive performance, as described in the main text and Tables S1–S5, rather than on the in-sample fits shown here. $R^2$ values for these fits are provided in the accompanying R Notebook HTML files.

**[DOWNLOAD]**

https://github.com/tksmiki/microscale_nonrandomness/releases/latest

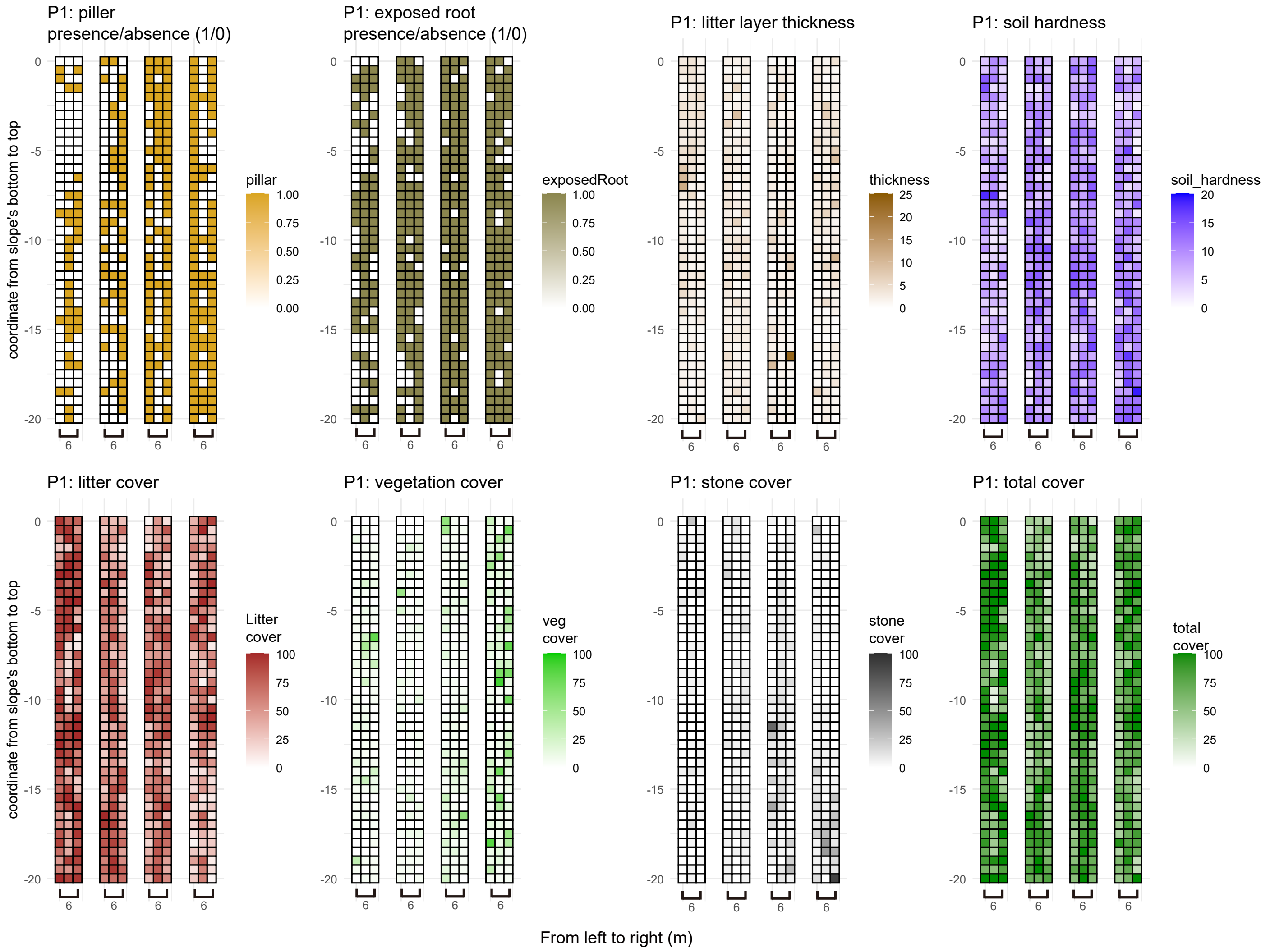

P1: piller presence/absence (1/0)
P1: exposed root presence/absence (1/0)
P1: litter layer thickness
P1: soil hardness
P1: litter cover
P1: vegetation cover
P1: stone cover
P1: total cover
coordinate from slope's bottom to top
From left to right (m)
pillar
exposedRoot
thickness
soil_hardness
Litter cover
veg cover
stone cover
total cover

**Figure S5. Eight microscale soil surface properties.** Spatial distributions of eight microscale soil surface properties that potentially characterize local soil conditions, including erosion risk (Miura *et al.*, 2015; Pellant *et al.*, 2020). Total cover (% of the observation window) is the sum of litter, vegetation, and stone cover. For dimensionality reduction by PCA, we included all eight variables, including total cover, despite its dependence on the three component cover variables.

#### SI References


Abe H., Kume T., Hyodo F., Oyamada M. & Katayama A. (2024a). Soil erosion under forest hampers beech growth: Impacts of understory vegetation degradation by sika deer. *CATENA* **234**, 107559. https://doi.org/10.1016/j.catena.2023.107559

Abe H., Kume T. & Katayama A. (2024b). Reduction in forest carbon stocks by sika deer-induced stand structural alterations. *Forest Ecology and Management* **562**, 121938. https://doi.org/10.1016/j.foreco.2024.121938

Andersen R.A., Berges J.A., Harrison P.J. & Watanabe M.M. (2005). Recipes for Freshwater and Seawater Media. pp. 429–538. Elsevier.

Bass C., Puinean A.M., Zimmer C.T., Denholm I., Field L.M., Foster S.P., *et al.* (2014). The evolution of insecticide resistance in the peach potato aphid, *Myzus persicae*. *Insect Biochemistry and Molecular Biology* **51**, 41–51. https://doi.org/10.1016/j.ibmb.2014.05.003

Beltrà A., Wäckers F.L., Nedvěd O. & Pekas A. (2018). Predation rate and performance of three ladybirds against the green peach aphid Myzus persicae in sweet pepper. *Entomologia Experimentalis et Applicata* **166**, 491–499. https://doi.org/10.1111/eea.12691

Frenken T., Alacid E., Berger S.A., Bourne E.C., Gerphagnon M., Grossart H.-P., *et al.* (2017). Integrating chytrid fungal parasites into plankton ecology: research gaps and needs: Research needs in plankton chytridiomycosis. *Environmental Microbiology* **19**, 3802–3822. https://doi.org/10.1111/1462-2920.13827

Guillard R.R.L. & Lorenzen C.J. (1972). YELLOW-GREEN ALGAE WITH CHLOROPHYLLIDE C. *Journal of Phycology* **8**, 10. https://doi.org/10.1111/j.0022-3646.1972.00010.x

Harada K., Ang Meng Ann J. & Suzuki M. (2020). Legacy effects of sika deer overpopulation on ground vegetation and soil physical properties. *Forest Ecology and Management* **474**, 118346. https://doi.org/10.1016/j.foreco.2020.118346

Ishihara M., Ishida K., Ida H., Itoh A., Enoki T., Ohkubo T., *et al.* (2010). An introduction to forest permanent plot data at core and subcore sites of the forest and grassland survey of the monitoring sites 1000 project. *Japanese Journal of Ecology* (in Japanese) **60**

Kagami M., Helmsing N.R. & van Donk E. (2011). Parasitic chytrids could promote copepod survival by mediating material transfer from inedible diatoms. *Hydrobiologia* **659**, 49–54. https://doi.org/10.1007/s10750-010-0274-z

Miki T., Takimoto G. & Kagami M. (2011). Roles of parasitic fungi in aquatic food webs: a theoretical approach: A food web model with parasitic fungi. *Freshwater Biology* **56**, 1173–1183. https://doi.org/10.1111/j.1365-2427.2010.02562.x

Miura S., Ugawa S., Yoshinaga S. & Keizo Hirai T.Y. (2015). Floor Cover Percentage Determines Splash Erosion in Chamaecyparis obtusa Forests. *Soil Science Society of America Journal* **79**, 1782–1791. https://doi.org/10.2136/sssaj2015.05.0171

Pellant M., Shaver P.L., Pyke D.A., Herrick J.E., Lepak N., Riegel G., *et al.* (2020). *Interpreting Indicators of Rangeland Health, Version 5*. U.S. Department of the Interior, Bureau of Land Management, National Operations Center, Denver, CO.

Seto K., Van Den Wyngaert S., Degawa Y. & Kagami M. (2020). Taxonomic revision of the genus *Zygorhizidium* : *Zygorhizidiales* and *Zygophlyctidales ord. nov.* ( *Chytridiomycetes* , *Chytridiomycota* ). *Fungal Systematics and Evolution* **5**, 17–38. https://doi.org/10.3114/fuse.2020.05.02

Smith V.H., McBride R.C., Shurin J.B., Bever J.D., Crews T.E. & Tilman G.D. (2015). Crop diversification can contribute to disease risk control in sustainable biofuels production. *Frontiers in Ecology and the Environment* **13**, 561–567. https://doi.org/10.1890/150094

Utsumi S., Nakamura M. & Ohgushi T. (2009). Community consequences of herbivore-induced bottom-up trophic cascades: the importance of resource heterogeneity: Induced resource heterogeneity for community. *Journal of Animal Ecology* **78**, 953–963. https://doi.org/10.1111/j.1365-2656.2009.01570.x

Widin S.L., Billings K.M., McGowen J. & Cardinale B.J. (2022). Biodiversity and disease risk in an algal biofuel system: An experimental test in outdoor ponds using a before-after-control-impact (BACI) design. *PLOS ONE* **17**, e0267674. https://doi.org/10.1371/journal.pone.0267674

Yoneya K., Miki T. & Katayama N. (2023a). Plant volatiles and priority effects interactively determined initial community assembly of arthropods on multiple willow species. *Ecology and Evolution* **13**, e10270. https://doi.org/10.1002/ece3.10270

Yoneya K., Miki T. & Takabayashi J. (2023b). Initial herbivory and exposure to herbivory-induced volatiles enhance arthropod species richness by diversifying community assemblages. *Frontiers in Ecology and Evolution* **10**, 1031664. https://doi.org/10.3389/fevo.2022.1031664
